# Cnidarian pharyngeal nervous system illustrates prebilaterian neurosecretory regulation of feeding

**DOI:** 10.1101/2023.04.05.535675

**Authors:** Ryo Nakamura, Ryotaro Nakamura, Hiroshi Watanabe

## Abstract

Neurosecretory brain centers exist as part of the central nervous system of many bilaterian animals and play pivotal roles in the control of feeding and metabolism. In the animal evolution, the nutrition regulation appears to have been essential for the early heterotrophic metazoan ancestor, but the underlying evolutionary processes remain unclear. We found that the cnidarian *Nematostella vectensis* develops an oral/pharyngeal nervous system that shares a core signature with bilaterian neurosecretory brain centers comprising *Orthopedia* and neuropeptide RFamide-positive neurons, and that is essential for feeding regulation. Our data suggest that prior to the bilaterian evolution, the *Orthopedia*/RFamide-positive neural assembly was already functioning as a prototype of the neurosecretory brain center for the feeding regulation that could be the driving force of neural centralization.

## Introduction

The evolutionary origin of the central nervous system (CNS) remains controversial^1–8^. Comparative analyses of a wide range of bilaterian lineages have shown considerable similarities and certain differences in the expression patterns of genes associated with the development and patterning of the neuroectoderm^1, 6–8^. One of the obstacles in pursuing the evolutionary origin of the CNS is the paucity of data focusing on the role of CNS patterning genes in the neurodevelopment, especially for phylogenetically informative lineages, due to the difficulty in analyzing gene functions in these non-model species. Further, the lack of data on comparable functional properties of neural structures hinders our understanding of the biological significance of the spatial arrangement of neural tissues and reconstruction of the key evolutionary processes underlying neural centralization. Members of the phylum Cnidaria, the closest sister group of bilaterians, are informative metazoan lineages for the understanding of the earliest assembly processes of the nervous system. Although cnidarians have a diffuse nervous system that lacks distinct neural tissue, their genomes possess many genes involved in the patterning and functioning of the bilaterian CNS. Still, the function of these genes remains largely unknown. Therefore, developmental and physiological comparisons between pre- and post-centralized nervous systems have been difficult^5, 8, 9^. The fundamental questions of how and why neural centralization came into play in the early evolution remains unanswered.

Here we report evolutionary conserved genetic and functional features between cnidarian oral/pharyngeal nervous system (PhNS) and bilaterian neurosecretory brain centers. The neurons comprising the PhNS in *N. vectensis* showed high expression of transcription factor *Orthopedia* (*Otp*) and neuropeptide RFamide, as well as genes for neuronal differentiation and chemical neurotransmission, which characterize bilaterian neurosecretory brain centers.

*Otp* knockdown resulted in downregulation of the PhNS genes, and the PhNS-deficient polyps showed reduced feeding ability, which was ameliorated by the activation of RFamide signaling. The results presented here suggest that the *Otp*/RFamide-positive neural subsystem was developed in functional association with mouth/pharynx as a primary neurosecretory center for feeding control before the divergence of the cnidarian and bilaterian lineages. Our findings also indicate that early neural centralization may have evolved primarily to regulate feeding-related responses.

## Results

### Cnidarian oral neurons share key molecules with bilaterian neurosecretory brain centers

In this study, we have focused on analyzing the developmental mechanisms and functions of the neural subsystem at the oral region of cnidarian *Nematostella vectensis*, where a number of neural genes including transcription factors (TFs) and neuropeptides are abundantly expressed^9–13^. Homologs of TFs involved in CNS patterning in Bilateria exist also in cnidarian genomes, and these homologs exhibit regionalized expression patterns along their oral-aboral and directive body axes (Fig. 1a; Supplementary Fig. 1)^2, 3, 5, 6, 9^. Among these homologs, our whole-mount *in situ* hybridization (WISH) analyses showed that the expression of *Otp* is specifically restricted to the oral side of the pharynx of *N. vectensis* (Fig. 1a), which is in agreement with previous study^11^. *Otp* is a well-conserved Paired (PRD)-class homeodomain containing TF found in Cnidaria and Bilateria (Supplementary Fig. 2; Supplementary Table 1), and expressed in the forebrain of annelids, insects, and vertebrates^14–21^. In vertebrates, *Otp* regulates the development of the hypothalamus^15, 16, 19^, which is a neurosecretory brain center that expresses a series of neuropeptides^19, 22^ to control pleiotropic homeostatic and behavioral functions^19^. Recent studies in *Platynereis dumerilii* (Annelida) and *Drosophila melanogaster* (Arthropoda) have revealed that some hypothalamus-related TFs and neuropeptides such as RFamide, are expressed in both species in the medial head/brain regions^17, 18, 21, 23–25^. These homologous molecular characteristics suggest that the neurosecretory brain center was already functional in the last common bilaterian ancestor. In cnidarians, peptide-expressing neurons develop into various neuronal types with specific spatial patterns. Among them, RFamide-expressing neurons localized to the oral/pharyngeal region have been widely observed in various cnidarian species and stages of their lifecycles (Fig. 1b)^9, 10, 12, 26, 27^. Although the specialized neural arrangement found around the mouth of cnidarians has long intrigued researchers, its relevance to the bilaterian CNS remained unclear.

**Fig. 1.**
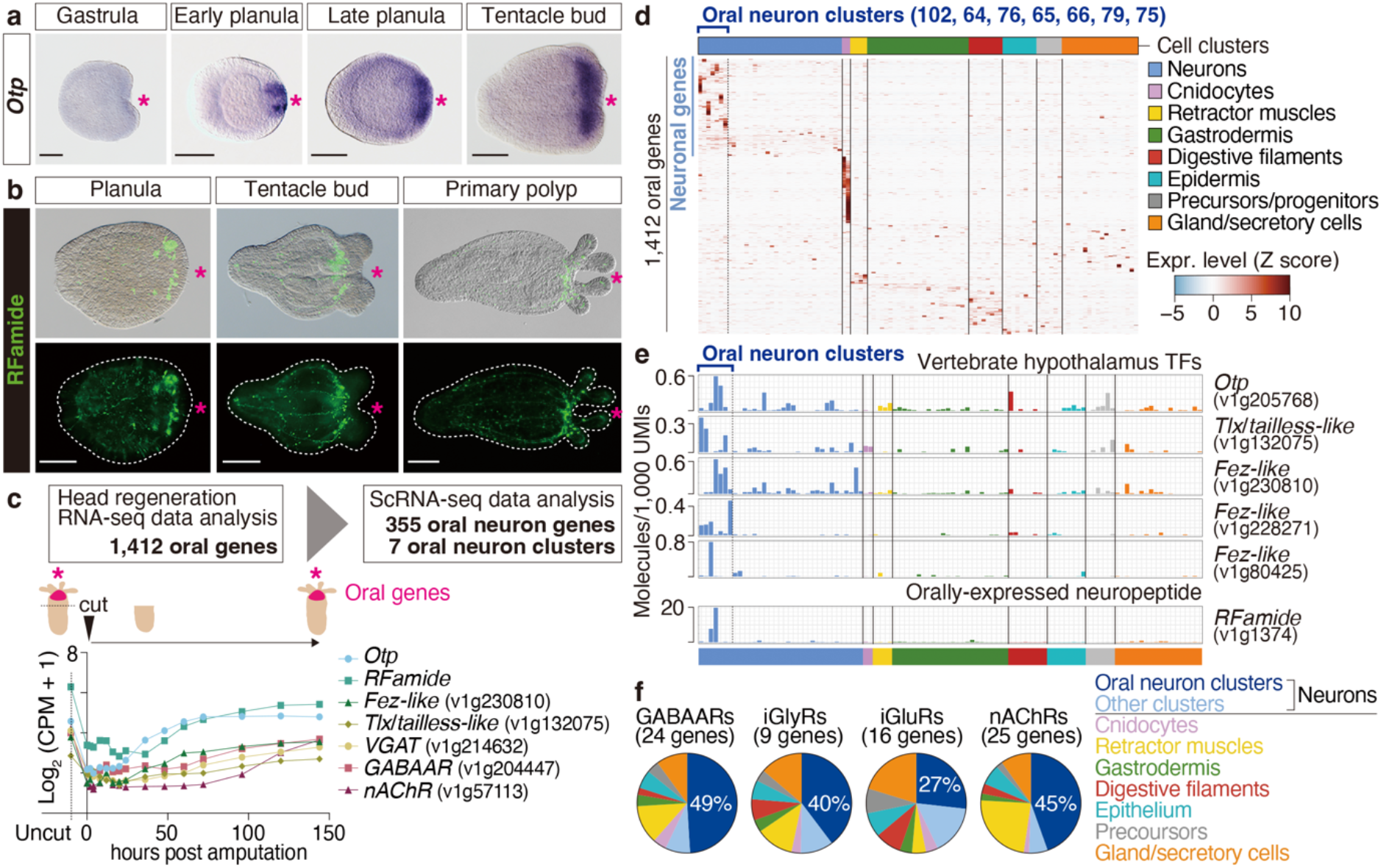
Characteristics of the *N. vectensis* pharyngeal nervous system (PhNS). (**a**) WISH analysis of *Otp* during *N. vectensis* development. (**b**) Immunohistochemical staining of RFamide from planula to primary polyp stages. (**c**) Scheme to identify oral neuron genes by retrieving *N. vectensis* adult RNA-seq data^28, 29^. The line plot shows the expression patterns of representative oral genes during head regeneration^29^. CPM, counts per million. (**d**) Distribution of the 1,412 identified oral genes in the adult cell clusters^28^. Gene expression levels for each cell cluster, determined from the z-scores of the log2 transformed values of the contigs, are displayed as colors ranging from blue to red (as shown in the key). The colors in the top indicate cell type of each cluster. (**e**) Expression of hypothalamus TF homologs and *RFamide* in adult *N. vectensis* across cell clusters^28^. The bars are colored according to the cell types shown in **d**. (**f**) Expressed cell types of putative ionotropic receptors for GABA (GABAAR), Gly (iGlyR), Glu (iGluR), and nicotinic ACh (nAChR) in adult *N. vectensis*^28^. The expression ratio for each cell type was calculated based on molecules/1,000 unique molecule identifiers (UMIs) reads per cell cluster (% of expression). Total % in oral neuron clusters are shown in the pie charts. Scale bars in **a**, **b**, 100 µm. The magenta asterisks in **a**–**c** indicate the oral or blastoporal side.

To determine the similarities between the oral assembly of the cnidarian neural subsystem and the neurosecretory brain centers of bilaterians, we examined the genetic and physiological features of neural subpopulations enriched in the mouth/head region at the cellular level using transcriptome datasets of genes expressed during body regeneration and at the single-cell level in *N. vectensis* which have recently been published^28, 29^. We carried out a comprehensive *in silico* survey of genes that are upregulated during regeneration of the oral tissue (head)^29^, identifying 1,412 genes as the orally-enriched genes (Fig. 1c). Combining these results with single-cell RNA-seq (scRNA-seq)^28^ data revealed that 36% of these genes were specifically expressed in the neuron clusters (Fig. 1d; Supplementary Fig. 3a). Among them, seven neuron clusters, metacell number 102, 64, 76, 65, 66, 79, and 75, were shown to express *Otp*, *Tlx*/*tailless*, *Fez*, and *RFamide* neuropeptide (Fig. 1e; Supplementary Fig. 3b, c); these genes are also enriched in the hypothalamus of vertebrates and neurosecretory brain centers of insects^14–16, 19, 22–25, 30^. Furthermore, in addition to these core genes, some chemical neurotransmitter transporters and receptors, including ionotropic receptors for gamma amino butyric acid (GABA), glycine (Gly), glutamate (Glu), and nicotinic receptors for acetylcholine (ACh)^13^, also showed abundant expression in these seven oral neuron clusters (Fig. 1f; Supplementary Fig. 3d). The transmitter complexity is reminiscent of the neurophysiological properties of hypothalamic neurons, which utilize chemical neurotransmitters in addition to peptide transmitters^31–33^. In this study, the neuronal populations that develop in the oral/pharyngeal region of cnidarians characterized by a distinct gene signature, will be referred to as the pharyngeal nervous system (PhNS).

### *Otp* regulates development of the cnidarian PhNS

To compare the genetic programs regulating the development of the cnidarian PhNS and bilaterian neurosecretory brain centers, we analyzed the function of *Otp* in the cnidarian PhNS development. Transfection of short interfering RNAs (siRNAs) against *Otp* significantly decreased *Otp* expression (Fig. 2a). In *N. vectensis* larvae (4 days post-fertilization [dpf]) transfected with siRNA1, which showed the highest *Otp* knockdown (KD) efficiency, we found 44 upregulated differentially expressed genes (DEGs) and 193 downregulated DEGs by RNA-seq analysis (Fig. 2b, c). Of the 193 downregulated DEGs, 19% were genes involved in gene transcription and 8% were related to neural signaling and transmission (ion channel subunits/transporters and receptors for chemical neurotransmitters and neuropeptides signaling molecules) which contained the *RFamide* gene (Fig. 2d). We confirmed downregulation of the neuronal DEGs (*RFamide*, transporters and receptors for GABA, Glu, and ACh) using quantitative polymerase chain reaction (qPCR) analysis (Fig. 2e). However, other genes expressed in the oral neuron clusters, such as *Tlx*/*tailless* and *Fez*, were not affected by *Otp* KD, suggesting that these genes act upstream of *Otp* or that their expression is regulated by a mechanism independent of *Otp*. The effect of *Otp* KD was less pronounced on *Elav* (Fig. 2e), which is widely expressed in endodermal neurons^34^, suggesting that *Otp* has a specific function in PhNS development, rather than being broadly involved in the neurodevelopment of the pervasive nerve net. Consistent with this, neurons expressing mature RFamide peptides were less detectable in the PhNS of *Otp* KD larvae, whereas the RFamide-positive nerve plexus remained intact (Fig. 2f, g). While *Otp* is mainly expressed in neurons (Fig. 1e), its function may not be essential for the gene regulatory network demarcation of blastoporal tissues because the expression levels of *Forkhead* (*FoxA*) and *Brachyury* were unaffected by *Otp* KD (Fig. 2e).

**Fig. 2.**
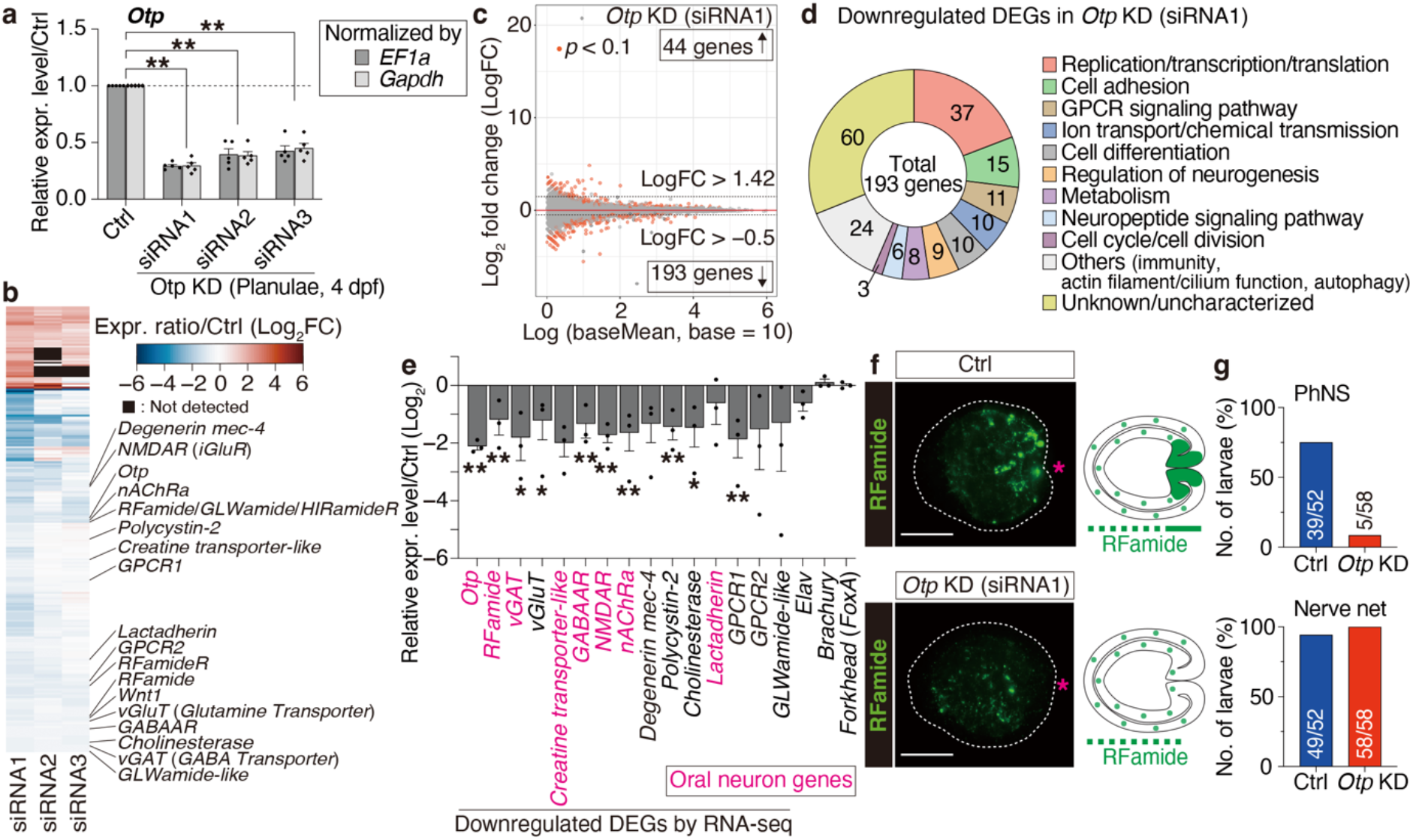
Effect of *Otp* knockdown on PhNS development. (**a**) Expression levels of the *Otp* mRNA in siRNA-transfected larvae assessed by qPCR. (N = 5). (**b**) Heatmap comparing expression levels of downregulated/upregulated genes among the three *Otp* knockdown (KD) conditions. Relative expression levels are colored from blue to red according to the scale. (**c**) MA plot of the gene expressions in the *Otp* KD (siRNA1) versus control larvae at 4 dpf. Orange points indicate genes with *P* < 0.1. (N = 5). (**d**) Functional categorization of downregulated DEGs in *Otp* KD (siRNA1) larvae. (**e**) qPCR assay of the expression levels of selected downregulated DEGs, *Elav*, *Brachury*, and *Forkhead* in *Otp* KD (siRNA1) larvae at 4 dpf, normalized to *Ef1a*. The oral neuron genes identified by *in silico* survey^28, 29^ are indicated in magenta. (N = 3). (**f**) Immunohistochemical staining of RFamide peptides in the control and *Otp* KD (siRNA1) larvae at 4 dpf. Representative distribution patterns of RFamide-positive neurons are illustrated on the right side. Scale bars, 100 µm. The magenta asterisks indicate the oral side. (**g**) Number of larvae (4 dpf) possessing a PhNS and/or nerve net in the control and *Otp* KD groups based on the RFamide immunostaining data. (N = 4). The error bars in **a**, **e** represent the mean ± standard error (*, *P* < 0.05; **, *P* < 0.01).

Our detailed expression and functional analyses of *Otp* in *N. vectensis* demonstrated that the genetic characteristics and developmental mechanisms of the PhNS are regulated differently from other neural subsystems, even in the less-assembled cnidarian nerve net^35^, and illustrated their similarity to the bilaterian neurosecretory brain centers. This newly detected similarity suggests that an *Otp*-dependent neuroendocrine system functionally deployed, at least in part, in the common ancestor of Cnidaria and Bilateria.

### The PhNS is essential for feeding-related behavior

What was the specific role played by this evolutionarily conserved *Otp*-dependent neurosecretory system of cnidarian PhNS? To gain insights into this question, we examined the function of *Otp*/RFamide-positive neurons of the PhNS by analyzing the behavior of *N. vectensis Otp* KD primary polyps. Although the KD effect of transient siRNA transfection on gene expression levels was attenuated at the polyp stage (Supplementary Fig. 4), a partial but significant level of inhibition on the development of RFamide-positive neurons of the PhNS was still detectable (Supplementary Fig. 5). We found out that in the *Otp* KD polyps at 6–7 dpf, the ingestion of prey rotifers (*Branchionus plicatilis*) was significantly decreased (Fig. 3a, b). Because the development of RFamide-positive neurons in the PhNS is dependent on *Otp* (Fig. 2e, f), the ingestion defect caused by *Otp* KD may have resulted from the attenuation of RFamide function in the PhNS. In fact, the feeding defect in the *Otp* KD polyps was recovered by the administration of synthetic mature RFamide peptide in a dose dependent-manner (Fig. 3b). The requirement of RFamide peptide in feeding was also confirmed by the decreased rotifer ingestion in *RFamide* KD polyps (Fig. 3c). We next investigated whether the role of the *Otp*/RFamide system in the PhNS is specific to local actions associated with the pharynx, or whether it also controls systemic responses concomitant with feeding. Spontaneous peristaltic movements along the trunk of *N. vectensis* polyps are known to be controlled by neural activity^13^. We found that peristaltic contractions were suppressed during feeding until the rotifers passed through the pharynx (Fig. 3d, e; Supplementary Fig. 6), suggesting that PhNS activity negatively regulates peristaltic contractions. In congruence with this, the PhNS developmental defects resulted in an increase of spontaneous peristaltic contraction frequency, which was suppressed by RFamide administration (Fig. 3f, g). These results indicate that *Otp*/RFamide-positive PhNS neurons not only control the ingestion process of the mouth/pharynx in close proximity to the PhNS but are also involved in the systemic responses associated with food intake (Fig. 3h).

**Fig. 3.**
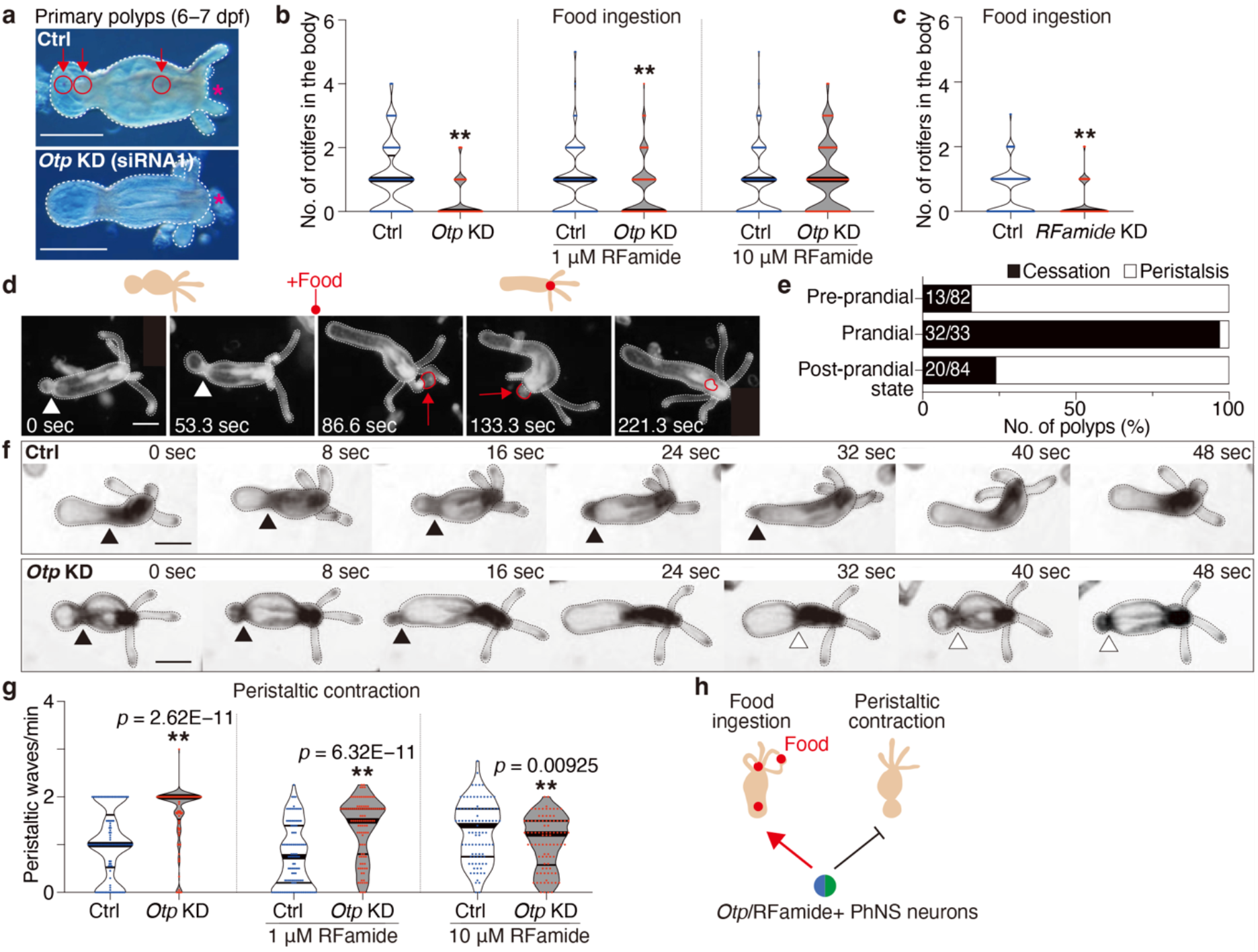
Role of *Otp*/RFamide-positive PhNS neurons in feeding behavior. (**a**) Representative pictures of control and *Otp* KD primary polyps after feeding with rotifers. The circles and arrows indicate the positions of the rotifers. The magenta asterisks indicate the oral side. (**b**) Counts of rotifers ingested in the control and *Otp* KD polyps at 6–7 dpf, treated with *N. vectensis* medium (N = 4) or either 1 or 10 μM synthetic RFamide peptide (N = 3). (**c**) Counts of rotifers ingested in the control and *RFamide* KD polyps at 6 dpf (N = 4). (**d**) Image series of a wild-type polyp at 6 dpf during feeding. Arrowheads indicate the contraction sites; red circles and arrows indicate the positions of rotifers. (**e**) The presence or absence of peristaltic movement from pre-to post-prandial states in the control polyps. (**f**) Images showing peristaltic waves in the control and *Otp* KD polyps. Black and white arrowheads indicate contraction sites of the first and second waves, respectively. (**g**) Frequencies of peristaltic waves per minute in the control and *Otp* KD polyps treated with *N. vectensis* medium (N = 4) or either 1 or 10 μM synthetic RFamide peptide (N = 3), respectively. (**h)** Putative function of *Otp*/RFamide-positive PhNS neurons. Scale bars in **a**, **d**, **f**, 250 µm. The black asterisks in **b**, **c**, **g** denote statistical significance with the Student’s t-test (**, *P* < 0.01).

## Discussion

In many marine invertebrates with simple body plans, neural cell assemblages have been widely observed in the proximity of oral and pharyngeal tissues^8–10, 12, 17, 20, 26, 27, 36–40^.

*Otp*/RFamide-positive cells show oral-regionalized expression patterns across species, especially in invertebrates, with these cells developing around the mouth and/or developing mouth regions (Fig. 4)^10–12, 17, 20, 36, 41^. Although these neuroanatomical arrangements observed in modern species do not necessarily indicate the typical morphological trait of ancestral neural assemblies, the fact that these neural configurations are broadly associated with the pharynx in marine invertebrates may provide insights into the functional aspects of what triggered the initial neural assembly process of the CNS. It is reasonable to assume that the primordial neural subsystem began to function to permit the local regulation of food ingestion, and was then expanded its role to regulate behavior and metabolism associated with nutrition intake.

**Fig. 4.**
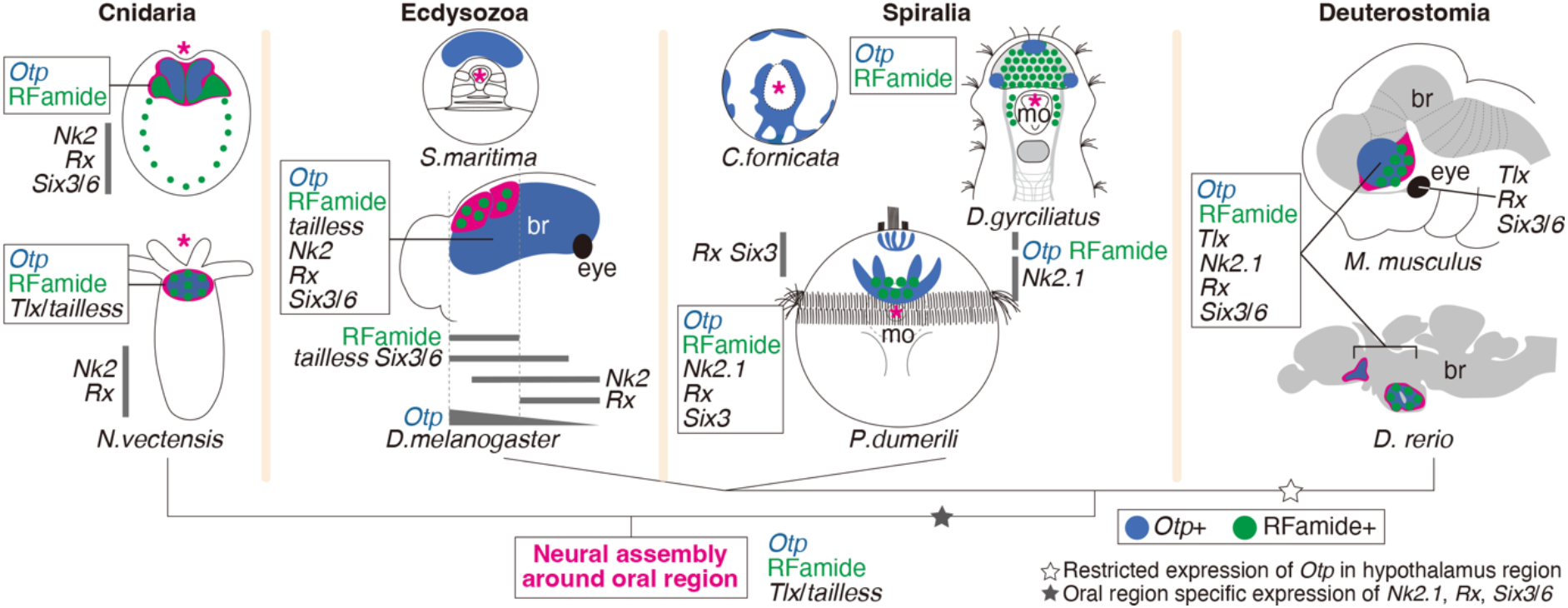
Evolutionary continuity of *Otp*/RFamide-positive neural assemblies among Cnidaria and Bilateria. Schematic drawings the expression patterns of *Otp*, RFamide, and homologs of bilaterian neurosecretory brain center TFs. *Otp* and RFamide expression are shown in blue and green, respectively. Cnidarian PhNS and known neurosecretory brain centers of insect and vertebrate (hypothalamus) are shown in magenta. The magenta asterisks indicate the oral side or mouth/developing mouth regions. br, brain; mo, mouth opening. The schemes are based on the following data: Cnidaria, *N. vectensis* (planula and polyp), compiled from this study and previous works^10–12, 45^; Ecdysozoa, *Strigamia maritima* (embryonic larvae, st 4.3)^36^ and *Drosophila melanogaster* (embryos)^14, 23–25^; Spiralia, *Crepidula fornicata* (elongation and early organogenesis stages)^41^, *Dinophilus gyrociliatus* (adult)^20^, and *Platynereis dumerilii* (trochophore larvae)^17, 18, 21^; Deuterostomia, *Mus musculus* (embryos)^14, 15, 19, 23, 25, 44^ and Danio rerio (adult)^16, 17, 19, 22, 25^.

The involvement of RFamide-family peptides in feeding regulation in vertebrates has been reported^42, 43^. In addition to *Otp*, *Tlx*/*tailless*, and *Fez* mentioned above, other TFs such as *Nk2.1*, *Rx*, and *Six3*/*6* also function in the hypothalamus/forebrain regions where RFamide peptides are expressed^17, 22, 23, 44^. Our data indicates that the PhNS of *N. vectensis* contains cells expressing *Tlx*/*tailless* and *Fez* (Fig. 1e, Fig. 4; Supplementary Fig. 1, Supplementary Fig. 3c). On the contrary, *Nk2*, *Rx*, and *Six3*/*6* genes showed aboral expression in Cnidaria (Supplementary Fig. 1)^11, 45^. To understand the evolutionary significance of the difference in arrangement of these genes in Cnidaria from Bilateria, their functions in cnidarian neural development will need to be determined.

In this work, we have provided experimental evidence of the evolutionary conservation of a neural subsystem associated with feeding, one of the most basic animal behaviors. Our findings in *N. vectensis* imply that some behavioral and metabolic control functions orchestrated by peptide-expressing neurosecretory brain centers in the CNS may have been functionally unitized by specific cell clusters in the diffused nervous system prior to its structural centralization. This provides a new evolutionary perspective on functional features that have been overlooked in the traditional approach of reconstructing the evolutionary origin of the CNS based on anatomically distinguishable neural structures. Previous studies have shown that cnidarians have several neural compartments with genetically distinct characteristics^12, 13, 34, 46^. Specific neural activities associated with distinct behavioral responses have recently been reported: in the freshwater polyp Hydra^47, 48^, which has a simple diffuse nervous system, and jellyfish^49^, which have developed distinct neural architectures, such as nerve rings. These findings lend further credence to the idea of functional compartmentalization in the pre-centralized nervous systems. Although it is possible that β-catenin and Bmp, which are important for the induction and patterning of the CNS in bilaterians, are involved in the development of the PhNS^12^, our understanding of the relationship between the functional compartments in the diffuse nervous system of cnidarians and the functional/structural compartments in the bilaterian CNS is still in its fancy. We believe that experimental verifications regarding whether cnidarian orthologs of neuroectoderm patterning genes in bilaterians are actually linked to the development/patterning of the cnidarian nervous system will reveal the initial state of the CNS and its evolutionary process in relation to the body axis.

## Methods

### Culture of N. vectensis

*N. vectensis* (Cnidaria) were cultured as described previously^12, 50^. Briefly, adult polyps were kept in *N. vectensis* medium (salinity of one-third of artificial seawater [35 g L^−1^, pH 7.5–8.0; SeaLife Marine Tech, Nihonkaisui Co., Ltd., Tokyo, Japan]) at 18°C in dark and fed twice a week with freshly hatched artemia. Spawning was induced by exposure to light overnight (O/N) at 26°C. After fertilization, embryos were cultured in *N. vectensis* medium at room temperature.

### Whole mount *in situ* hybridization (WISH) analysis

To generate digoxigenin (DIG)-labelled antisense probes, the target gene fragments were amplified from a cDNA library. The primer sets for PCR cloning are listed in Supplementary Data 1. PCR products were cloned into vector pGEM-T (Promega, Madison, WI), linearized, and *in vitro* transcribed with sp6 or T7 RNA polymerase (Roche, Basel, Switzerland)) according to the insert direction. The sequences of *Gsx*, *Otx*, and *Gsx*, *Nk2* homologs were taken from Finnerty et al. (2003)^51^, Mazza et al. (2007)^52^, and Steinmetz *et al*. (2017)^53^, respectively.

WISH was performed as described previously^12, 54^ with the following modifications: specimens were fixed with 4% (w/v) paraformaldehyde (PFA)/phosphate buffered saline (PBS, pH 7.4) at 4°C for 1 hour (gastrulae [2 dpf]) or O/N (early [3 dpf]/late [4 dpf] planulae and tentacle bud stage [5 dpf]). Following fixation, the specimens were washed with PBST (PBS, 0.1% Tween 20) and then transferred into methanol. The specimens were stored at −20 °C. The signal was visualized with BM Purple (Roche).

### Immunohistochemistry analysis of RFamide

Immunohistochemistry (IHC) was performed as described previously^12, 50^ with the following modifications: for the primary polyps (5 dpf), the animals were anesthetized in 1.5% MgCl_2_· 6H_2_0 in *N. vectensis* medium for 10 minutes to allow them to settle down prior to fixation; larvae and polyps were fixed with Zamboni’s fixative (15% picric acid, 2% PFA, 0.1 M phosphate buffer [pH 7.2]) O/N at 4°C; fixed specimens were transferred into methanol then washed with PBST (PBS, 0.1% Tween 20); after permeabilization in PBST (PBS, 0.1% Triton X-100), the specimens were incubated in blocking solution (1% bovine serum albumin [BSA], 5% Normal goat serum, 0.1% sodium azide, 0.2% PBST [Tween 20]) O/N at 4 °C. Anti-RFamide antibody (1:1,000 in the blocking solution) generated based on identification by mass spectrometry analyses^50^ and Alexa488-conjugated anti-rabbit antibody (1:500; Jackson ImmunoResearch Laboratories, Inc., West Grove, PA) were used as the primary and secondary antibodies, respectively. All antibodies were incubated O/N at 4°C.

### Gene expression profiling analysis

*In silico* analyses were performed by utilizing published scRNA-seq^28^ and head regeneration RNA-seq^29^ data of *N. vectensis* adults. All genes were annotated based on reciprocal Basic Local Alignment Search Tool (BLAST) searches (blastp, version 2.11.0)^55^ using *D. rerio* proteins as queries. The protein sequences of nAChRs and some of the GABAARs of *N. vectensis* were taken from published studies^13, 46^. Putative G-protein coupled receptors (GPCRs) for the neuropeptides were identified using a prediction method described previously^50^.

### Phylogenetic sequence analysis

Sequences of PRD-class homeodomain genes were collected from GenBank^56^ or the Joint Genome Institute (JGI) genome database for *N. vectensis* (v1.0)^57^. Putative *Otp* sequences were identified by reciprocal BLAST searches with blastp^55^ with an e-value cut-off of 0.05 against the NCBInr^58^ and JGI genome databases^57^ using *D. rerio Otp* proteins as queries. HMMER (version 3.2.1)^59^ was used to determine the sequences of homeodomain and *otp*, *aristaless*, and *rax* (OAR) motif regions. For phylogenetic tree generation, all 61 protein sequences were aligned by *ClustalW*^60^ in MEGA software (version X)^61^. The aligned sequences are listed in Supplementary Data 2. For each gene, partition models^62^ were utilized to specify and calculate the best substitution models according to each domain/motif sequence of the gene: part 1, 1–294 bp of the aligned sequences, VT+F+R3; part 2, 295–361 bp (homeodomain region), LG+G4; part 3, 362–875 bp (region including the OAR motif), VT+F+G4. The phylogenetic tree was built with PhyML 3.0^63^ with maximum-likelihood (ML) analysis in IQ-TREE (version 1.6.12)^64^. A total of 10,000 ultrafast bootstraps were calculated^65^. *Shox*, *Arx*, and *Unc-4* served as outgroups to root the tree. Tree visualization was performed by FigTree software (v1.4.4, http://tree.bio.ed.ac.uk/software/figtree/).

### siRNA-mediated KD in *N. vectensis*

To KD genes, siRNAs were electroporated into *N. vectensis* eggs based on methods we recently developed^50, 66^ and as described previously^67^ with the following modifications: the eggs were suspended in *N. vectensis* medium with 8% Ficoll before electroporation. After electroporation, the eggs were incubated for 2 hours with sperm-containing *N. vectensis* medium for fertilization. Fertilized eggs were transferred into fresh *N. vectensis* medium; for *Otp* KD, the eggs were electroporated with siRNA at 250 ng µL^-1^. KD efficiencies were assessed by qPCR analyses; for *RFamide* KD, we performed the same procedures as with the *Otp* KDs, except that the final concentration of siRNA was at 100 ng µL^-1^. The sequences of the siRNAs were as follows: *Otp* siRNA1 (5’-CGAUCAGGACGACGACAGAdTdT-3’), *Otp* siRNA2 (5’-CGGAUGUGUUCAUGCGAGAdTdT-3’), *Otp* siRNA3 (5’-GUCAGCAAUCGCAGCAGAAdTdT-3’), *RFamide* (5’-GCUUGGAAUCCUAAUUCAAdTdT-3’) and control (5’-GCAACACGCAGAGUCGUAAdTdT-3’).

### RNA-seq

siRNA-mediated KD larvae at 4 dpf were collected for RNA-seq. Total RNA was extracted using the RNeasy kit (QIAGEN, Hilden, Germany), and poly(A) was purified. cDNA libraries were sequenced on the NovaSeq 6000 SP system (paired end; Illumina, San Diego, CA). Five biological replicates were processed for each experimental condition (control and *Otp* KD [siRNA1–3] groups, 20 samples). We obtained a total of 1.95 B reads, with a minimum of 67.7 M reads per sample. The quality of the reads was assessed with FastQC (http://www.bioinformatics.babraham.ac.uk/projects/fastqc/). Reads were mapped to the reference genome of *N. vectensis* downloaded from https://metazoa.ensembl.org/info/data/ftp/index.html and annotated using Hisat2^68^. Transcript abundances were quantified using featureCounts^69^. Library preparation and sequencing were conducted by the OIST Sequencing Section.

### Differentially expressed gene (DEG) screening

Differential expression analysis in the *Otp* KD and control groups was performed using the DESeq2 method^70^. Transcripts with missing or total read counts of less than 10 for all samples were excluded before carrying out the analysis. The values calculated by DESeq2 are listed in Supplementary Data 3. DEGs with an adjusted *p* value of less than 0.1 were sorted using a lo_g2_ fold change cut-off of −0.5 for downregulated genes and +1.42 for upregulated genes. Putative genes that were derived from bacterial genomes were excluded. Downregulated DEGs in *Otp* KD (siRNA1) were annotated by BLAST alignment to the NCBInr database^58^ of *D. rerio*, *M. musculus* and *Homo sapiens* with an e-value cut-off of 0.05. All putative homologs were confirmed by reciprocal blastp^55^ using *D. rerio* proteins as queries (Supplementary Data 4).

### qPCR analysis

Total RNA was isolated from *N. vectensis* using the RNeasy kit (QIAGEN) and reverse transcribed into cDNA. qPCR analysis was performed on a StepOne Plus™ Real-Time PCR System (Thermo Fisher Scientific, Waltham, MA) using the PowerUp SYBR® Green Master Mix (Thermo Fisher Scientific). The expression levels of analyzed genes were normalized to *Ef1a* and/or *Gapdh* using the ΔΔCt method. The qPCR primer sequences are listed in Supplementary Data 5.

### Analysis for *N. vectensis* feeding behaviors

To assess the feeding behaviors of *N. vectensis*, siRNA-mediated KDs of *Otp* (siRNA1) and *RFamide* were performed as described above and primary polyps at 6–7 dpf were assayed. The polyps were placed in a BSA-coated slide chamber 1 minute before the start of recording. Then, 100 µL of rotifers (*Branchionus plicatilis sp. complex*) suspended in *N. vectensis* medium was added 1 minute after the start of recording. The concentration of rotifers was adjusted to the same level for each experiment. For RFamide peptide administration, the peptide solution was added from two opposite sides of the slide chamber and then mixed carefully using a pipette. Five minutes after the administration of the peptide solution, 100 µL of the rotifer-containing solution was added. The final concentration of the peptide solution was either 1 or 10 µM in *N. vectensis* medium. The sequence of the synthesized *N. vectensis* RFamide peptide was as follows: PyrGRFGREDQGRFamide. The number of rotifers that had been ingested by the polyps was counted 9 minutes after adding the rotifers. The number of peristaltic waves in the polyps was counted before adding the rotifers. The RFamide peptide was synthesized by Scrum Inc. (Tokyo, Japan). Rotifers were purchased form Nikkai Center (Tokyo, Japan).

### Microscopy

The stained larvae in WISH and IHC analyses were imaged with a DS-L3-equipped ECLIPSE Ni-E fluorescence microscope (Nikon Corporation, Tokyo, Japan). Time-lapse recordings of the polyps during feeding behavior analysis were captured using an SZX16 stereo microscope (Olympus, Tokyo, Japan).

### Data analysis

Fluorescent microscope images were processed with NIS-Elements AR (Version 5.02.03, Nikon). Image data were edited using ImageJ/Fiji (version 2.1.0/1.53c). Numerical data were analyzed using R (version 4.2.2) or Microsoft Excel and visualized using RStudio (2022.07.2+576) or GraphPad Prism (version 9).

### Statistics and reproducibility

Statistical data analysis was performed using Microsoft Excel. The statistical significance for qPCR and feeding behavior analyses were determined using paired samples *t*-test (two-tailed distribution). The DEG analysis of RNA-seq data was conducted using DESeq2^70^ and *p* values were caluculated. Statistical results were indicated with asterisks as follows (\**p* < 0.05, \*\**p* < 0.01). The details of error bars for each experiment are annotated in figure legends. Multiple trials at least three times were performed for all experiments.

### Reporting Summary

Further information on research design is available in the Nature Portfolio Reporting Summary linked to this article.

### Data availability

RNA-seq data conducted in this study were deposited in DNA Data Bank of Japan Sequence Read Archive with the submission number DRA015632. The data generated or analyzed in the RNA-seq analysis are included in the Supplementary Data 3 and 4.

## Acknowledgments

We thank the OIST Sequencing and Scientific Computing & Data Analysis Sections for their support in RNA-seq; J. Higuchi and A. Tanimoto for maintaining the *N. vectensis* culture; C. Guzman and H. Wang for their help on retrieving scRNA-seq data; A. Joshi for his technical assistance on siRNA evaluation. We are extremely grateful to P. Barzaghi, M. Miyake, L. Sheloukhova, M. Dindo, and OIST science editor, M. Cheung for careful reading and improvement of the manuscript.

## Author contributions

Ryo N. performed most of the experiments and all data analyses. H.W. and Ryotaro N. performed WISH analyses. Ryo N. and H.W. designed this project and wrote the manuscript. All authors have approved the final manuscript.

## Additional information

<u>Supplementary Information (Word .docx file)</u>

This file contains supplementary figures and table.

**Supplementary Figures 1–6**

**Supplementary Table 1**

<u>Separate supplementary materials</u>

**Supplementary Data 1** (Excel .xlsx file)

List of primers used to amplify genes for WISH probes synthesis.

**Supplementary Data 2** (.txt file)

Aligned sequences of genes *Otp, Shox, Arx*, and *Unc-4* used for phylogenetic tree generation.

**Supplementary Data 3** (Excel .xlsx file)

List of gene IDs and calculated values by DESeq2 from the RNA-seq data in *Otp* KD larvae at 4 dpf (siRNA1–3).

**Supplementary Data 4** (Excel .xlsx file)

Annotated information of downregulated DEGs in *Otp* KD larvae (siRNA1).

**Supplementary Data 5** (Excel .xlsx file)

List of primers for qPCR analyses.

