## Supplementary information file for "Cnidarian pharyngeal nervous system illustrates prebilaterian neurosecretory regulation of feeding"

**1. Supplementary figures**

**Supplementary Fig. 1**

**Supplementary Fig. 2**

**Supplementary Fig. 3**

**Supplementary Fig. 4**

**Supplementary Fig. 5**

**Supplementary Fig. 6**

**2. Supplementary table**

**Supplementary Table 1**

### 1. Supplementary figures

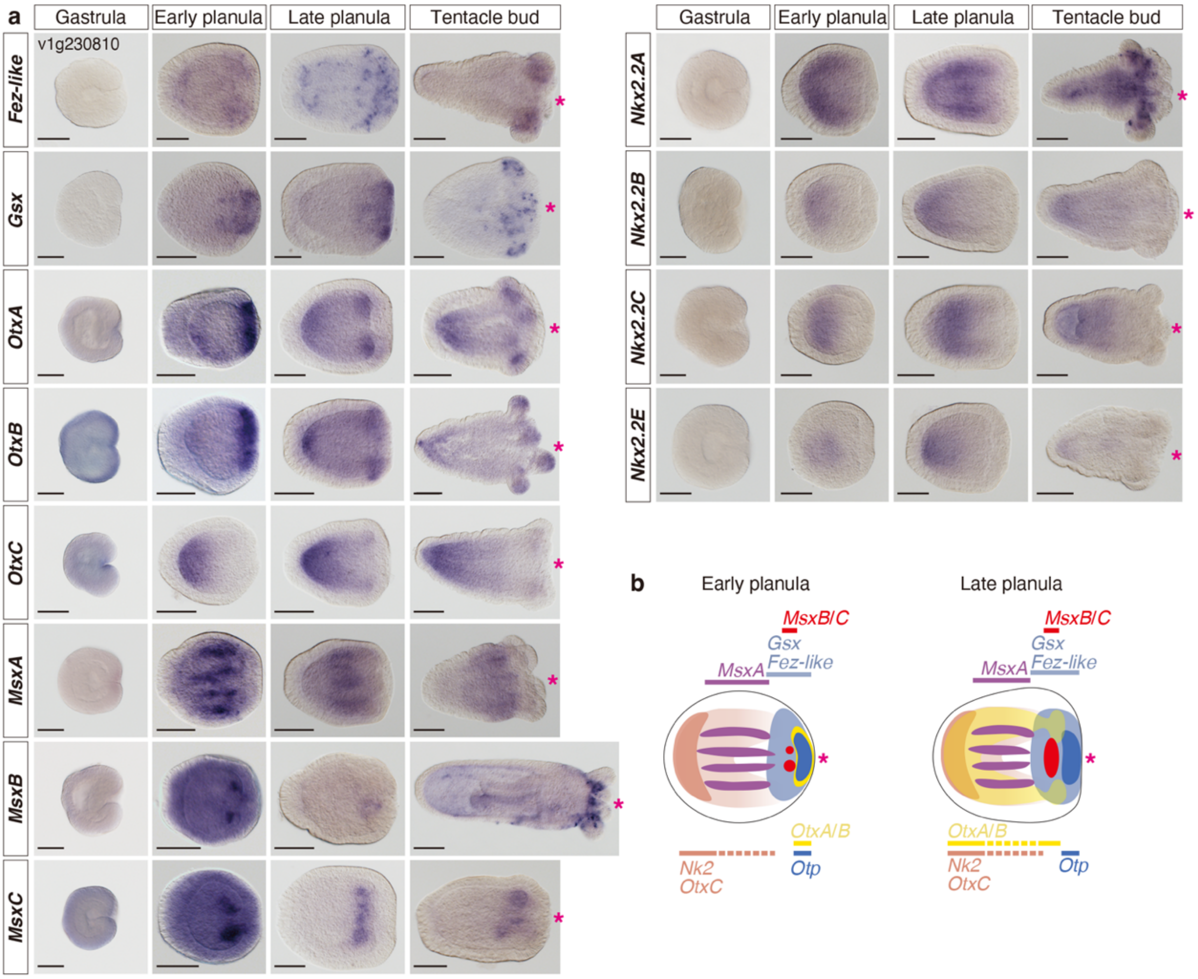


**Supplementary Fig. 1. Expression patterns of *N. vectensis* TFs involved in CNS patterning in Bilateria.** (**a**) WISH analysis for *Fez*, *Gsx*, *Otx*, *Msx*, and *Nk2* (*Nkx2*) genes from gastrula to tentacle bud stages. Scale bars, 100 µm. (**b**) Schematic diagrams showing expression patterns of *N. vectensis* TFs in early and late planula larvae. The magenta asterisks indicate the oral side.

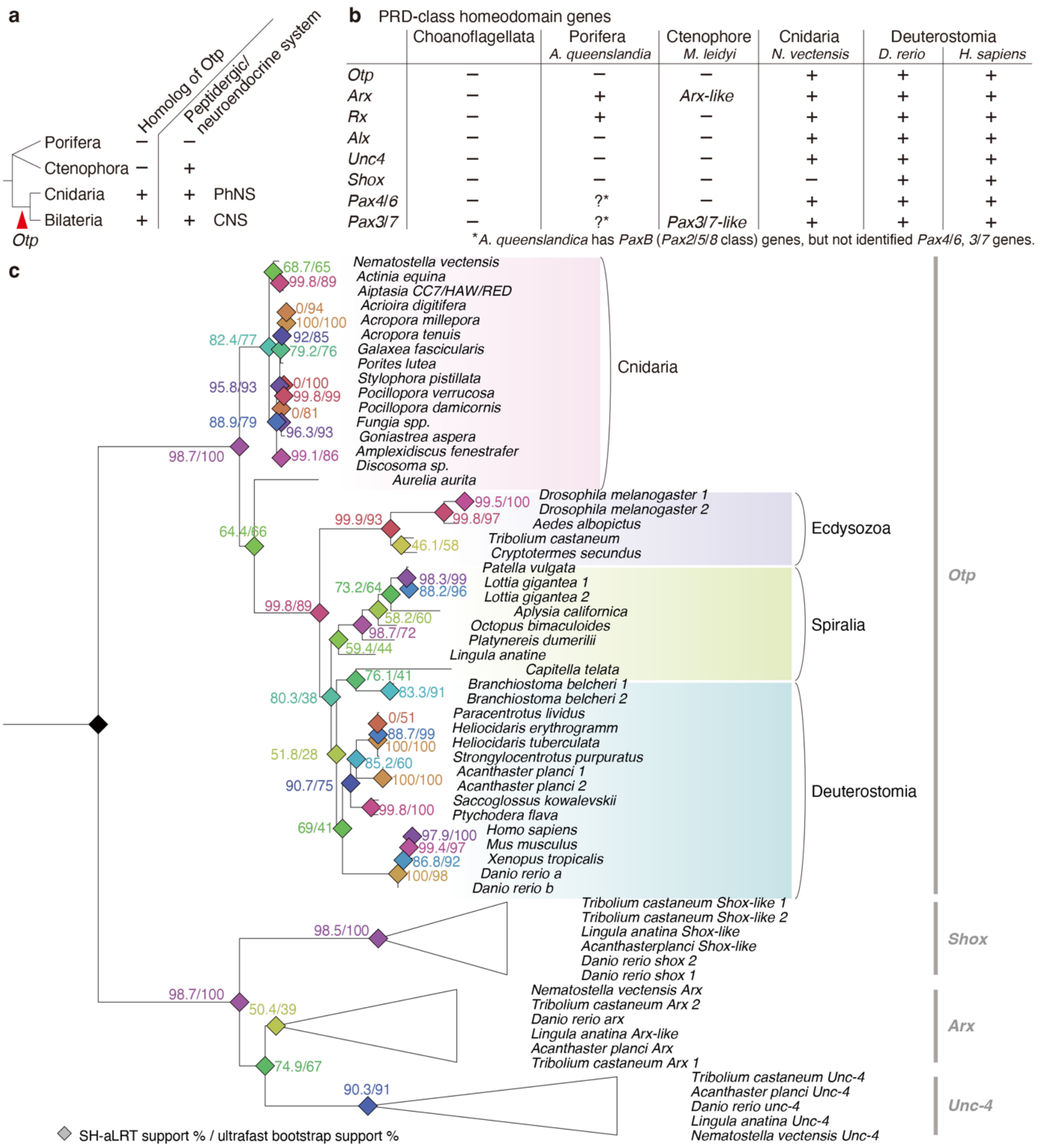


**Supplementary Fig. 2. Phylogenetic relationship of *Otp*.** (**a**) Schematic diagram showing the relationship of nervous system evolution and appearance of *Otp*. (**b**) PRD-class homeodomain proteins including *Otp* in Choanoflagellata, Porifera (*Amphimedon queenslandica*), Ctenophore (*Mnemiopsis leidyi*), Cnidaria (*N. vectensis*), and Deuterostomia (*D. rerio*, *H. sapiens*). Note: *Otp* is conserved in Cnidaria and Bilateria. (**c**) Phylogenetic tree of *Otp* genes in Cnidaria, Ecdysozoa, Spiralia, and Deuterostomia. SH-aLRT, Shimodaira-Hasegawa approximate likelihood ratio test.

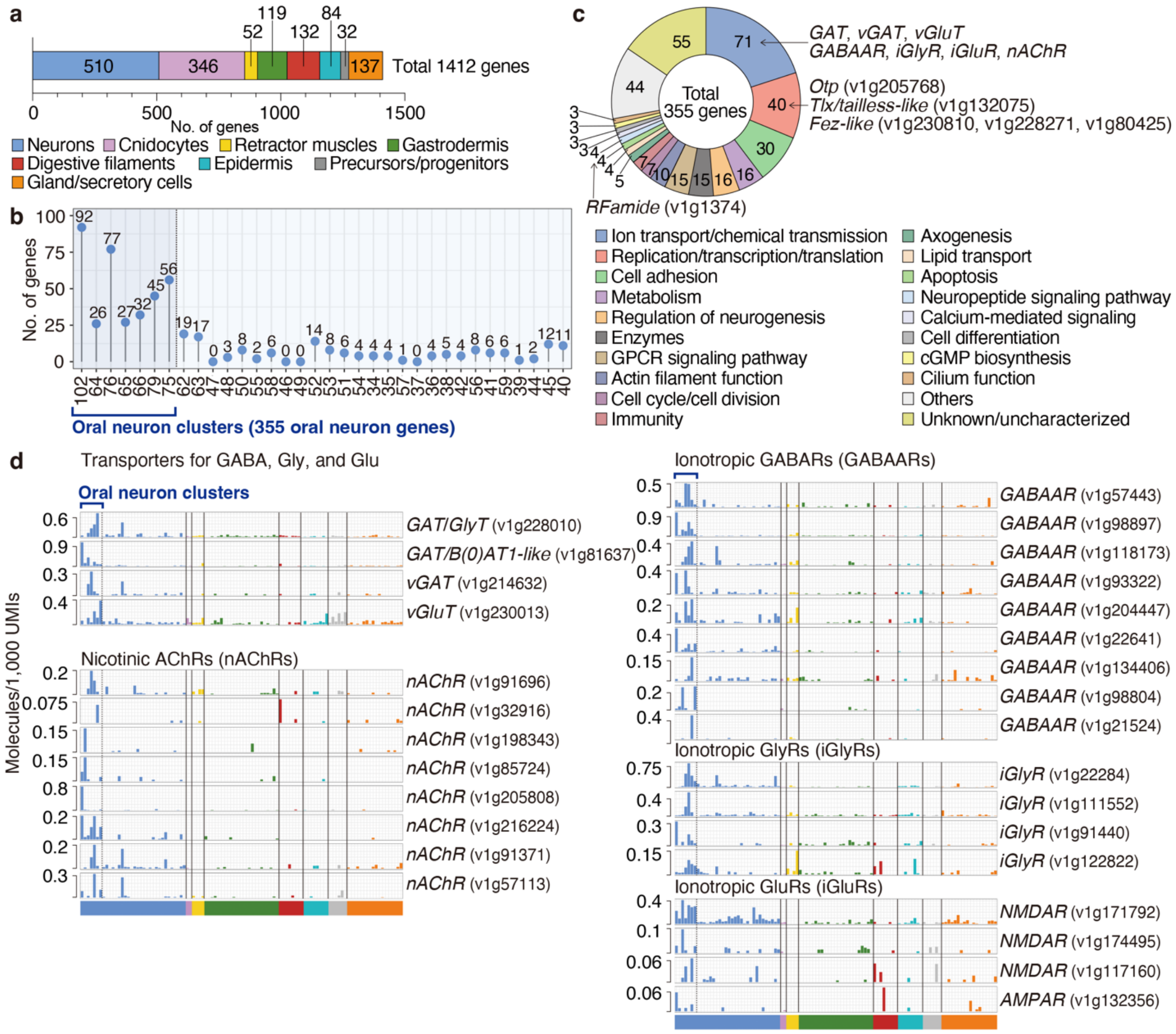


**Supplementary Fig. 3. Identification and characterization of *N. vectensis* oral neuron genes.** (**a**) Expressed cell types of the 1,412 identified oral genes in adult *N. vectensis*[28]. The cell type for each gene was determined based on the cell cluster with the highest expression. Note: 510 genes show the highest expression in neurons. (**b**) Distribution of the 510 genes showing highest expression in neurons across adult neuron clusters[28]. Seven neuron clusters showing particular enrichment of these oral genes were denoted as oral neuron clusters in this study. (**c**) Functional categorization of 355 oral neuron genes in the seven oral neuron clusters. (**d**) Expression (molecules/1,000 UMIs) of selected transporters and ionotropic receptors for chemical neurotransmitters in adult *N. vectensis*[28]. The bars are colored according to the cell types shown in **a**. GAT, GABA transporter; vGAT, vesicular GABA transporter; vGluT, vesicular glutamate transporter; GlyT, Glycine transporter; B(0)AT1,

System B(0) neutral amino acid transporter 1; NMDAR, N-methyl-D-aspartate receptor; AMPAR, α-amino-3-hydroxy-5-methyl-4-isoxazolepropionic acid receptor.

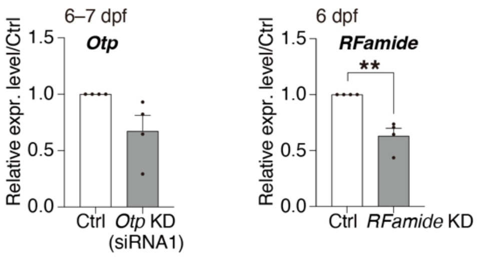


**Supplementary Fig. 4. Efficiency of siRNA-mediated KD in *N. vectensis* polyps.** Gene KD levels in primary polyps developed from siRNA-transfected eggs assessed by qPCR (N = 4). Error bars represent the mean ± standard error (**, $P < 0.01$).

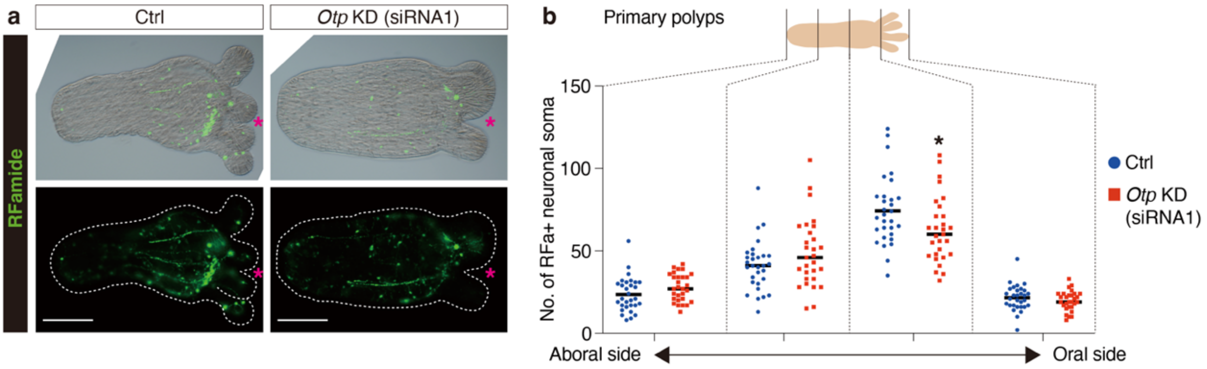


**Supplementary Fig. 5. Distribution of RFamide-positive neurons in *Otp* KD polyps.** (**a**) Immunohistochemical staining of RFamide in control and *Otp* KD early primary polyps (5 dpf). Scale bars, 100 µm. The magenta asterisks indicate the oral side. (**b**) Comparison of the RFamide-positive neuronal soma numbers along the oral-aboral axis in the control and *Otp* KD polyps (N = 3). The black asterisk denotes statistical significance using the Student's t-test (*, $P < 0.05$).

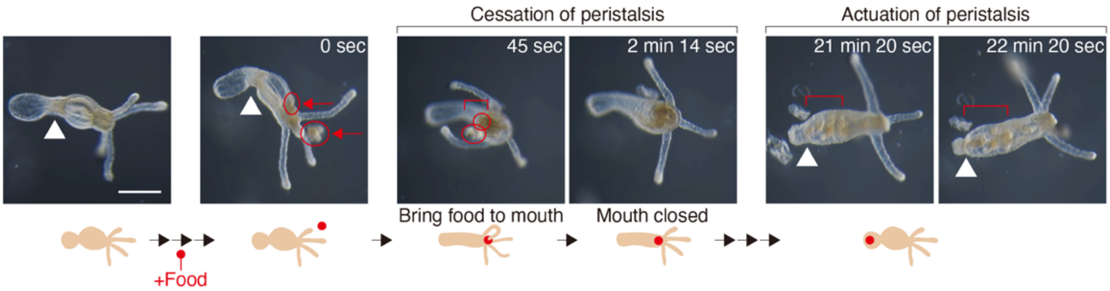


**Supplementary Fig. 6. Time-course of peristaltic movement during feeding.** Image series of a wild-type polyp at 6 dpf from pre- to post-prandial states. Arrowheads indicate the contraction positions; red circles, arrows, and square brackets indicate the positions of the rotifers. Note: peristaltic contraction was actuated again after the rotifers had passed through the pharynx. Scale bar, 250 µm.

### 2. Supplementary table

| Group | Phylum | Superphylum | Species name | No. of *Otp* |
|---|---|---|---|---|
| Amoedozoa | | | | 0 |
| Fungi | | | | 0 |
| Nuclearidae | | | | 0 |
| Filasterea | | | | 0 |
| Choanoflaggelate | | | solitary / colonial | 0 |
| Porifera | | | | 0 |
| Ctenophore | | | | 0 |
| Placozoa | | | *Trichoplax adhaerens* | 1 |
| Cnidaria | | Anthozoa | *Nematostella vectensis* | 1 |
| | | Anthozoa | *Acrioira digitifera* | 1 |
| | | Anthozoa | *Acropora tenuis* | 1 |
| | | Anthozoa | *Amplexidiscus fenestrafer* | 1 |
| | | Anthozoa | *Discosoma sp.* | 1 |
| | | Anthozoa | *Fungia spp.* | 1 |
| | | Anthozoa | *Galaxea fascicularis* | 1 |
| | | Anthozoa | *Goniastrea aspera* | 1 |
| | | Anthozoa | *Pocillopora damicornis* | 1 |
| | | Anthozoa | *Pocillopora verrucosa* | 1 |
| | | Anthozoa | *Porites lutea* | 1 |
| | | Anthozoa | *Actinia equina* | 1 |
| | | Anthozoa | *Aiptasia* | 1 |
| | | Scyphozoa | *Aurelia aurita* | 1 |
| | | Stylophora | *Stylophora pistillata* | 1 |
| | | Hydrozoa | *Hydra magnipapillata/vulgaris* | 1 |
| | | Hydrozoa | *Hydra viridissima* | 1 |
| | | Hydrozoa | *Cladonema pacificum* | 0 |
| | | Hydrozoa | *Clytia hemisphaerica* | 0 |
| Ecdysozoa | Arthropoda | Insecta | *Drosophila melanogaster* | 2 |
| | Arthropoda | Insecta | *Tribolium castaneum* | 1 |
| | Arthropoda | Insecta | *Aedes albopictus* | 1 |
| | Arthropoda | Insecta | *Clastoptera arborina* | 0 |
| | Arthropoda | Insecta | *Cryptotermes secundus* | 1 |
| | Priapulida | Priapulimorpha | *Priapulus caudatus* | 1 |
| Spiralia | Mollusca | Gastropoda | *Patella vulgata* | 1 |
| | Mollusca | Gastropoda | *Lottia gigantea* | 2 |
| | Mollusca | Gastropoda | *Aplysia californica* | 1 |
| | Mollusca | Cepharopoda | *Octopus bimaculoides* | 1 |
| | Brachipoda | Lingulata | *Lingula anatine* | 1 |
| | Brachipoda | Articulata | *Terebratalia transversa* | 1 |
| | Annelida | Polychaeta | *Platynereis dumerilii* | 1 |
| | Annelida | Polychaeta | *Capitella telata* | 1 |
| | Platyhelminthes | Turbellaria | *Schmidtea polychroa* | 1 |
| | Platyhelminthes | Cestoda | *Sparganum proliferum* | 1 |
| | Platyhelminthes | Rhabditophora | *Schistosoma mansoni* | 1 |
| Deuterostomia | Echinoderm | Asteroidea | *Acanthaster planci* | 2 |
| | Echinoderm | Echinoidea | *Strongylocentrotus purpuratus* | 1 |
| | Hemichordata | Enteropneusta | *Saccoglossus kowalevskii* | 1 |
| | Hemichordata | Enteropneusta | *Ptychodera flava* | 1 |
| | Chordata | Urchordata | *Ciona intestinalis* | 1 |
| | Chordata | Cephalochordata | *Branchiostoma belcheri* | 2 |
| | Chordata | Mammalia | *Homo sapiens* | 1 |
| | Chordata | Mammalia | *Mus musculus* | 1 |
| | Chordata | Amphibia | *Xenopus tropicalis* | 1 |
| | Chordata | Osteichthyes | *Danio rerio* | 2 |
| Xenacoemorpha | Xenacoelomorpha | Nemertodermatida | *Meara stichopi* | 1 |
| | | Acoela | *Convolutriloba longifissura* | 1 |

**Supplementary Table 1. List of *Otp* in basal metazoan animals.** The presence or absence and the number of *Otp* homologs in holozoans and metazoans.
